# Single-molecule Mechanical Analysis of Strand Invasion in Human Telomere DNA

**DOI:** 10.1101/2021.06.22.449520

**Authors:** Terren R. Chang, Xi Long, Shankar Shastry, Joseph W. Parks, Michael D. Stone

**Affiliations:** Department of Chemistry and Biochemistry, University of California, Santa Cruz, 1156 High St, Santa Cruz, CA 95064, USA; 10X Genomics, 6230 Stoneridge Mall Rd, Pleasanton, CA 94588; Invitae, 1400 16th St, San Francisco CA 94103, USA

## Abstract

Telomeres are essential chromosome end capping structures that safeguard the genome from dangerous DNA processing events. DNA strand invasion occurs during vital transactions at telomeres, including telomere length maintenance by the alternative lengthening of telomeres (ALT) pathway. During telomeric strand invasion, a single stranded guanine-rich (G-rich) DNA invades at a complimentary duplex telomere repeat sequence forming a displacement loop (D-loop) in which the displaced DNA consists of the same G-rich sequence as the invading single stranded DNA. Single stranded G-rich telomeric DNA readily folds into stable, compact, structures called G-quadruplexes (GQ) in vitro, and is anticipated to form within the context of a D-loop; however, evidence supporting this hypothesis is lacking. Here we report a magnetic tweezers assay that permits the controlled formation of telomeric D-loops (TDLs) within uninterrupted duplex human telomere DNA molecules of physiologically relevant lengths. Our results are consistent with a model wherein the displaced single stranded DNA of a TDL folds into a GQ. This study provides new insight into telomere structure and establishes a framework for development of novel therapeutics designed to target GQs at telomeres in cancer cells.

## INTRODUCTION

Telomeres safeguard the genome by distinguishing chromosomal termini from sites of DNA lesions which would otherwise elicit an unwanted DNA damage response resulting in chromosomal fusion, genomic instability, and often apoptosis ^1, 2^. The foundation of telomere structure begins with tandem hexameric guanine-rich (G-rich) repetitive DNA (GGTTAG in humans) ∼2 – 20 kilobases in length ^3, 4^ and terminates with a ∼50 – 300 nucleotide long single stranded G-rich 3’ overhang ^5^. Telomeres also act to buffer against the end replication problem, wherein chromosomes gradually shorten with each subsequent round of cell division ^6^. Replication-dependent telomere attrition can compromise the protective function of telomeres as well as lead to a loss of genetic information if left unaddressed ^7^. Therefore, continually dividing cells, including the majority of human cancers, must maintain telomere length to support an immortal phenotype ^8, 9^. A majority of proliferative cell types upregulate the specialized enzyme telomerase, which reverse transcribes telomeric DNA on to chromosomal termini using an RNA template that resides within the integral telomerase RNA subunit ^10-13^. However, many aggressive cancer subtypes employ a telomerase-independent mechanism for telomere maintenance termed alternative lengthening of telomeres (ALT) ^14^. In ALT cells, the 3’ single stranded DNA (ssDNA) overhang of one telomere base-pairs with the duplex region of another telomere, in a manner similar to early steps in homology directed repair ^15^. This telomeric strand invasion event forms a displacement loop (D-loop), where the single stranded G-rich 3’ overhang base-pairs with the C-rich strand of the invaded telomere, displacing the G-rich strand ^16-18^. The 3’ overhang can then be extended by a specialized DNA polymerase using the invaded telomere as a template ^19^, followed by C-strand fill in and nucleolytic processing to maintain the 3’ overhang ^20^.

Single stranded G-rich telomeric DNAs readily fold into compact structures called G-quadruplexes (GQ) *in vitro*, wherein guanine bases form G-quartets via both Watson-Crick and Hoogsteen base-pairing interactions to align in a plane while coordinating a monovalent cation at the center. Multiple G-quartets can in turn stack upon each other form a GQ ^21^. The stability of GQs is highly dependent on the identity of the monovalent cation, with a rank order of K^+^ > Na^+^ > Li^+^, in terms of degree of stabilization ^22^. Furthermore, small molecule ligands designed to target GQ structures elicit a phenotype in living cells, suggesting a possible regulatory role for these structures *in vivo* ^23^. Therefore, much effort has been put forth in identifying potential GQ forming sequences in the genome to expand the potential targets for these molecules to be used as therapeutics ^24^. In the current study, we report results from a single-molecule mechanical assay of DNA strand invasion at human telomeres. Using a magnetic tweezers system, uninterrupted duplex telomere DNA molecules as long as ten kilobases can be manipulated in order to impart precise degrees of tension and torque to the system. Strand invasion by single-stranded DNA oligonucleotides in solution can be monitored in real-time as a change in the overall extension of the telomere DNA duplex target molecule. To our knowledge, this assay represents the first to permit direct detection of telomeric D-loops (TDLs) at the single-molecule level. We find that conditions which disfavor GQ folding dramatically alter the properties of TDLs, suggesting a role for GQ folding within these important structures. Finally, this system provides an experimental framework for future singlemolecule studies of small molecule drugs and cellular machinery that may bind and alter GQ structure within a TDL.

## RESULTS

### Single molecule manipulation of long human telomere DNA molecules

The DNA molecules used in the present work consist of between nine to twelve kilobases of uninterrupted double-stranded telomeric DNA sequence. The telomere DNA molecule is flanked by biotin- or digoxigenin-modified DNA linker fragments used to immobilize the DNA tether between a streptavidin-coated magnetic bead and an anti-digoxigenin coated glass slide, respectively (Figure 1A and 1B). To generate these long, uninterrupted, telomere DNA tether molecules for singlemolecule analysis in our magnetic tweezers microscope, we perform a controlled DNA concatenation reaction seeded on the digoxigenin linker fragment using a 576 base pair telomere DNA fragment with compatible sticky ends generated by restriction endonuclease cleavage of the previously reported pRST5 DNA plasmid ^25^. Following multiple rounds of DNA ligation, the molecule is ultimately capped by ligation of the biotin-modified DNA linker fragment and gel purified to remove unwanted reaction side products and excess handle material (Figure 1A, see Methods for details of DNA molecule construction).

**Figure 1.**
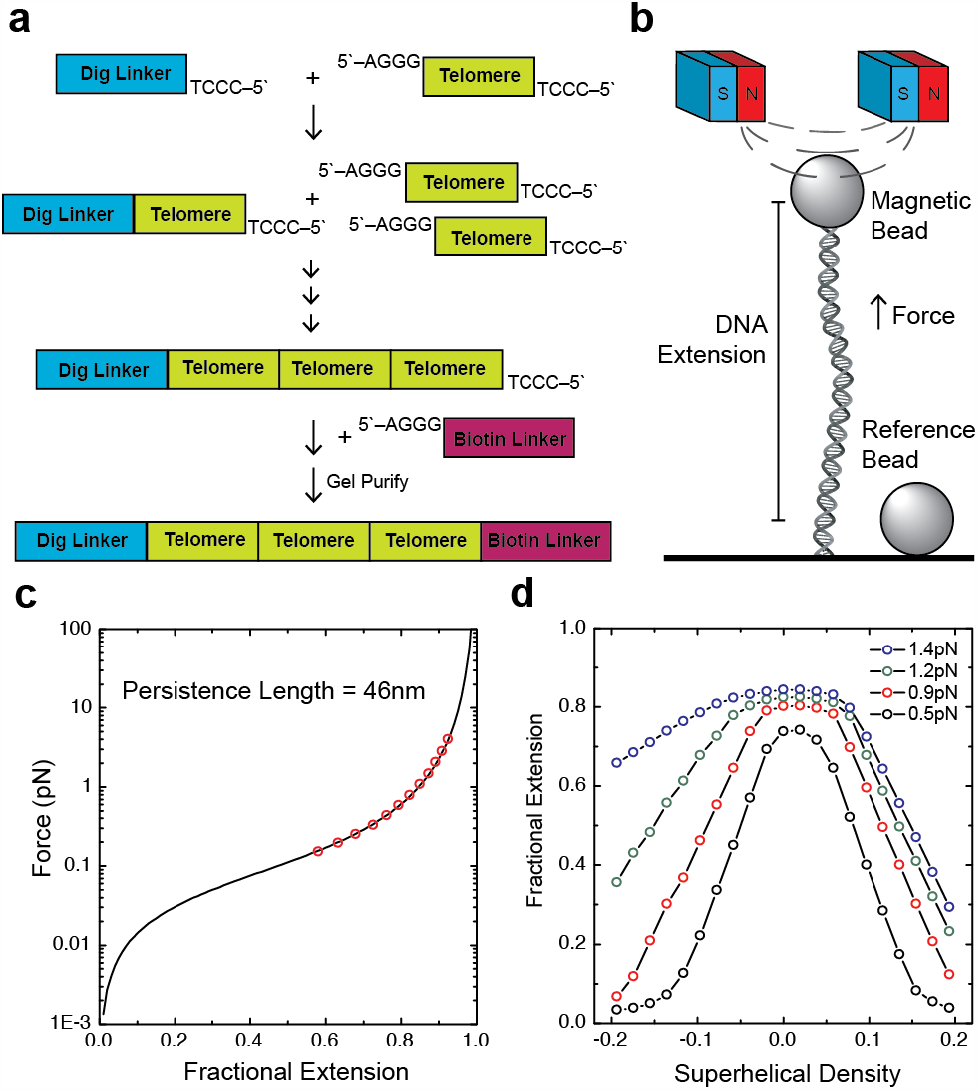
Mechanical properties of long duplex telomeric DNA. (a) Schematic of the construction of telomeric DNA molecules used in this study. (b) Schematic of the magnetic tweezers instrument. The vertical position of the magnets is adjusted to control the force exerted on the tethered DNA molecule. The magnets can also be rotated to apply torque. (c) Force extension curve of telomeric DNA. Data points are in red with the wormlike chain fit in black. (d) Rotation extension curves of telomeric DNA at various forces.

The elasticity of doubled stranded DNA is well described by the worm-like chain (WLC) polymer model ^26^ and is characterized by a bending persistence length ranging from ∼45-50 nanometers (nm), depending upon the ionic strength ^22, 27^. To test whether our telomere DNA tethers exhibit similar elastic properties, we performed force-extension analysis. Our results indicate that long double-stranded telomere DNA molecules exhibit canonical DNA elastic properties with an average persistence length of 46 +/- 4 nm under the conditions of our experiments (10 mM Tris pH 7.5, 150 mM KC_2_H_3_O_2_, 0.5 mg/mL BSA) (Figure 1C). Next, we analyzed the supercoiling response of our telomere DNA tethers by rotating the magnets held above a molecule of interest, which permits precise introduction of positive or negative superhelical strain into the system. DNA tether extension data is collected for a variety of supercoiling densities, given by the expression σ = ΔL_k_ / L_ko_, where σ is the supercoiling density, ΔL_k_ is the change in DNA linking number (ie. integer number of magnet rotations), and L_ko_ is the linking number of the DNA molecule in a topologically relaxed state (ie. the total number of DNA base pairs in the tether divided by the number of base pairs per helical turn of the double helix) (Figure 1D). Interestingly, we find that telomeric DNA more readily denatures in response to applied negative superhelical strain when compared to a non-telomeric control DNA (Figure S1), consistent with a recently reported study of force-induced denaturation of a non-telomeric GQ forming sequence ^28^.

### Real-time observation of DNA strand invasion in human telomere DNA

Having characterized the physical properties of the telomere DNA tethers, we next developed a DNA topology-based assay to directly measure telomere DNA strand invasion in real time (Figure 2A). The molecule is initially negatively supercoiled resulting in a decrease in extension. When the molecule is stretched to 0.9 pN of force, the negative superhelical density imparts torque on the molecule, which results in transient, local destabilization of the DNA double helix and facilitates strand invasion by a freely diffusing complementary DNA oligonucleotide from solution ^29^ (Figure 2A and 2B). In this assay, the negatively supercoiled telomere DNA tether represents a closed topological system. Therefore, the local DNA unwinding that must occur upon strand invasion imparts compensatory positive supercoiling into the molecule, which in turn cancels some of the preexisting negative supercoiling, resulting in a sudden increase in the DNA tether extension when held at constant force (Figure 2B, black arrowheads). To initially characterize strand invasion in our system we monitored the properties of a 42 nucleotide long single-stranded invading DNA molecule comprised of seven repeats of the C-rich telomere DNA strand sequence (Tel7C). Although this oligonucleotide is the complement of the physiologically relevant G-rich ssDNA telomere tail, it has the advantage of not folding into GQ structures and therefore simplified the interpretation of our initial experiments. In the presence of 10 nM Tel7C we observe discrete steps of increased extension as a function of time as expected (Figure 2B). Importantly, in the absence of complementary oligonucleotides in solution we do not observe any stable changes in the DNA tether extension (Figure S2). A collection of time trajectories as shown in Figure 2B are fit with a step finding algorithm ^30^ to extract the distributions of step sizes and waiting times between discrete stepping events (Figure S3). For the Tel7C invading strand the distribution of step sizes is 176 ± 6 nm. Once a DNA molecule buckles and begins to form plectonemic supercoils, subsequent supercoiling results in a linear change in the DNA tether extension, yielding an extension change of 50 nm per turn introduced under our experimental conditions (Figure 2C). Using this value, we would predict that invasion of the 42 nt long Tel7C molecule should remove ∼ 4 helical turns (42 bases / 10.5 base pairs per turn), which should result in 200 nm steps in the extension signal. The ∼10% lower change in extension we observe for the tel7C invasion events may be due to incomplete invasion of the entire oligonucleotide sequence. However, we do find that the mean step size does monotonically change with different length C-rich oligonucleotides (Tel3C and Tel15C, Figure 2D and Figure S4) as expected, supporting the conclusion that the discrete steps we observe in the real-time strand invasion trajectories correspond to individual oligonucleotides invading the target molecule and forming telomeric displacement loops (TDLs).

**Figure 2.**
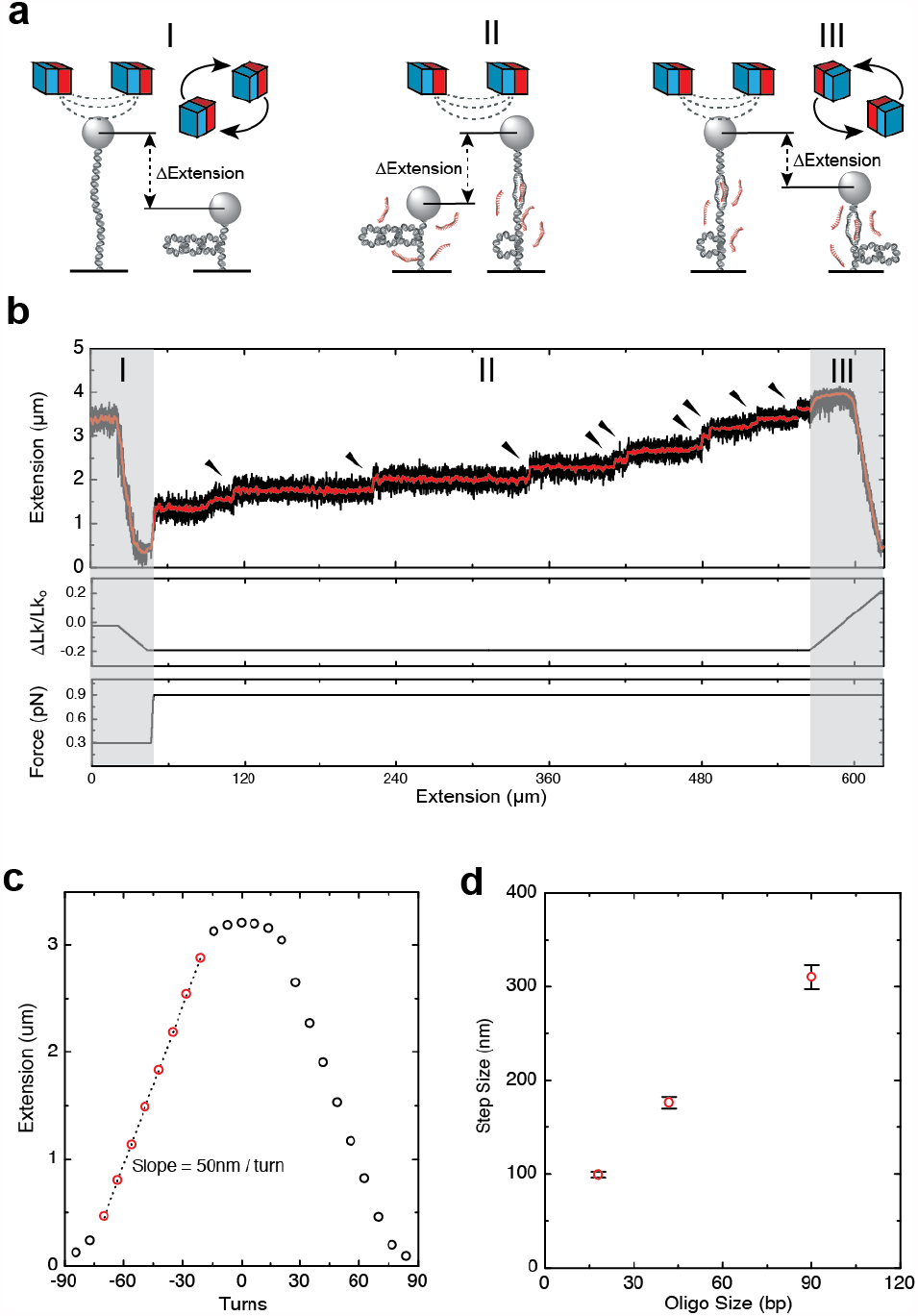
Real-time detection of telomeric D-loop formation by strand invasion. (a) Schematic of the magnetic tweezers experimental setup. Initially, a single telomere DNA molecule is tethered between a glass slide and a magnetic bead, followed by introduction of negative supercoils (I). Spontaneous strand invasion events are detected as discrete, stepwise increases in DNA tether extension (II) (black arrowheads in panel b). The molecule is then positively supercoiled while monitoring the extension (III). (b) Real-time trajectory of a Tel7C (CCCTAA)7 strand invasion experiment. Top panel indicates raw DNA extension signal (black) overlaid with a one second moving average (red). The middle panel shows the supercoiling density of the DNA tether throughout the experiment, expressed as ΔLk/Lk_o_. (c) Rotation extension curve at 0.9 pN. At this force, the rotation extension curve is symmetrical, with a linear response of tether extension to applied negative supercoiling as shown by the linear fit to the red symbols (dashed line) with a slope of 50 nm per turn introduced. (d) The observed change in extension upon strand invasion correlates with the length of the strand used for invasion. The calculated mean step size is plotted from experiments conducted with Tel3C, Tel7C, and Tel15C having 3, 7, and 15 C-rich repeats, respectively. Error bars are the standard error of the mean.

### Formation of telomeric displacement loops is torque dependent

To gain insight into the energetics of strand invasion for the tel7C strand we next monitored the torque-dependence of the rate of strand invasion by studying the dwell time distributions at a variety of force set points (Figure 3). In our system, the amount of torque (τ) applied locally to the DNA molecule can be approximated by the expression τ = (2BF)^1/2^, where B is the DNA bending persistence length and F is the applied stretching force ^31^. Interestingly, we find the kinetics of strand invasion to be exceedingly torque-dependent, which can be seen qualitatively by comparison of representative strand invasion trajectories collected at different stretching forces (Figure 3A). For each of the force set points, a distribution of waiting times between strand invasion events can be measured and the mean dwell time for each experiment is plotted as a function of the applied torque (Figure 3B). This torque-velocity relationship can be modeled using a simple transition state model, given by the expression *k*(τ) = *k*_*o*_exp^-2Πnτ^, where *k*_*o*_ is the rate constant for the strand invasion process in the absence of applied torque, and n is the characteristic number of base pairs unwound in the target DNA at the transition state for the strand invasion reaction ^32^. Fitting the torque-dependence of the kinetics of strand invasion with this simple model yields values of *k*_*o*_ = 3 × 10^−7^ *s*^−1^ and n = 9 base pairs (Figure 3B and 3C). This analysis indicates that only a small fraction of the 42 nt invading strand is required to base pair with the target DNA to create an effective toe hold for the formation of a D-loop, and in the absence of applied torque the formation of these structures occurs at a vanishingly slow rate.

**Figure 3.**
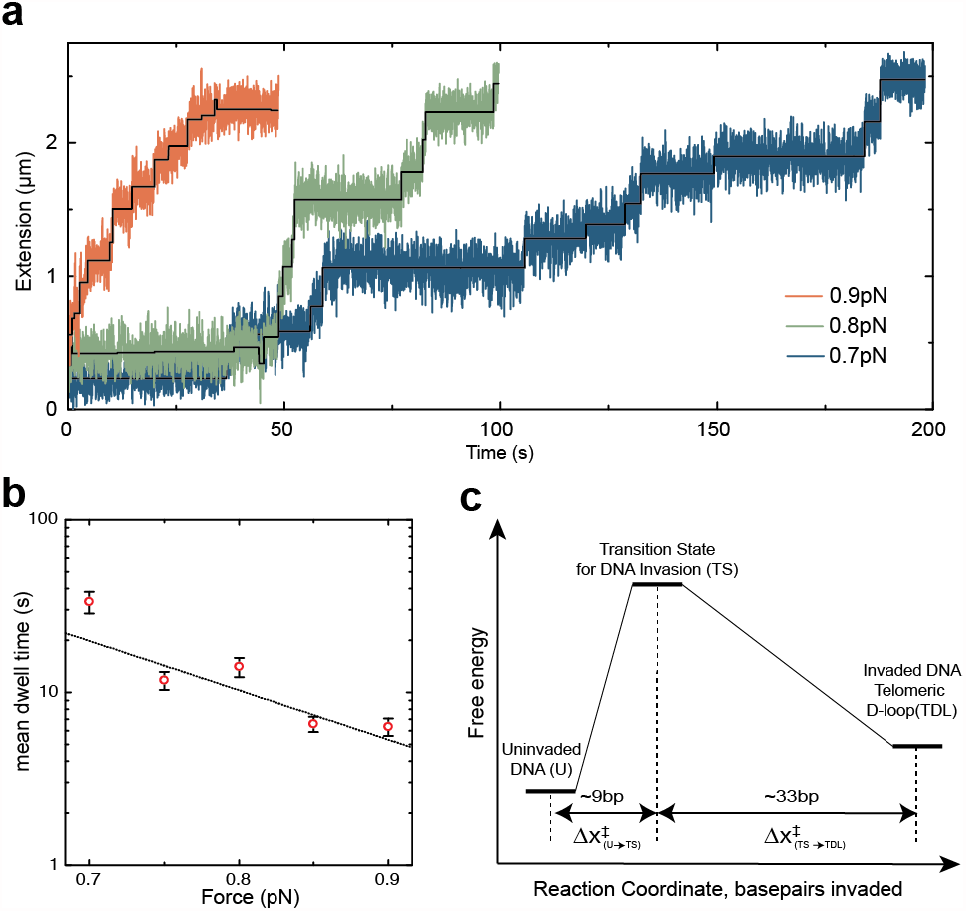
Kinetic analysis and energetics of telomeric strand invasion. (a) Time traces of 2*μ*M tel7C strand invasion at 0.7pN (blue), 0.8pN (green) and 0.9pN (red). Idealized trajectory derived by stepfitting algorithm shown in black. (b) Plot of mean dwell time between invasion events as a function of torque. Dotted line is a single exponential fit. Error bars are the standard error of the mean. (c) Free energy landscape of strand invasion reaction progressing from the initial uninvaded DNA (U) state to the invaded DNA telomeric displacement loop (TDL). Using a simple two-state kinetic model, the invasion process passes through a single high energy transition state (TS). From the exponential fit in Figure 3b, we find an ∼ 9 base pair separation between the initial uninvaded DNA (U) and the transition state (TS) along the reaction coordinate as the reaction progresses to complete the formation of a telomeric D-loop.

### G-rich telomere DNA exhibits complex structural dynamics during strand invasion

We next set out to investigate differences in the strand invasion dynamics observed for the C-rich tel7C strand as well as the complimentary G-rich seven-repeat telomere sequence (tel7G) (Figure 4). Comparison of strand invasion trajectories collected for the tel7C and tel7G invading strands reveal an obvious qualitative difference. Whereas the tel7C invasion trajectory primarily consists of a stepwise monotonic increase in the observed DNA tether extension as a function of time (Figure 4A, top panel), the process for the tel7G invading strand exhibits increased complexity characterized by a higher prevalence of backstepping (Figure 4A compared to Figure 4B). Despite this distinction, waiting time distributions for individual invasion events collected for both the Tel7C and Tel7G are well-fit by single exponential functions with similar decay constants (Figure S5). We note that invasion of the more physiologically relevant Tel7G strand generates a telomere D-loop wherein the displaced strand possesses the identical Grich telomere repeat sequence. Therefore, we reasoned that the increased complexity and backstepping observed in the Tel7G strand invasion trajectories may be due in part to the propensity of either the G-rich invading or displaced strand to form GQ structures. To investigate whether GQ folding dynamics within the invading strand are necessary for the observed complexity, we performed control experiments with a modified G-rich invading oligonucleotide in which the second base of each consecutive G-triplet was replaced with a 7-deazaguanine modified base. Invading oligonucleotides harboring this 7-deazaguanine modification cannot form the requisite Hoogsteen hydrogen bonds required to form a stable GQ structure ^33^. When using this 7-deazaguanine modified strand (tel7dG) we found that the frequency of backstepping in the invasion traces was comparable to that observed with the native tel7G sequence (Figure S6). Thus, the observed dynamics are not dependent upon GQ formation within the invading G-rich strand, suggesting that the sequence of the displaced strand folds into a GQ structure within a TDL formed upon invasion of the tel7G sequence.

**Figure 4.**
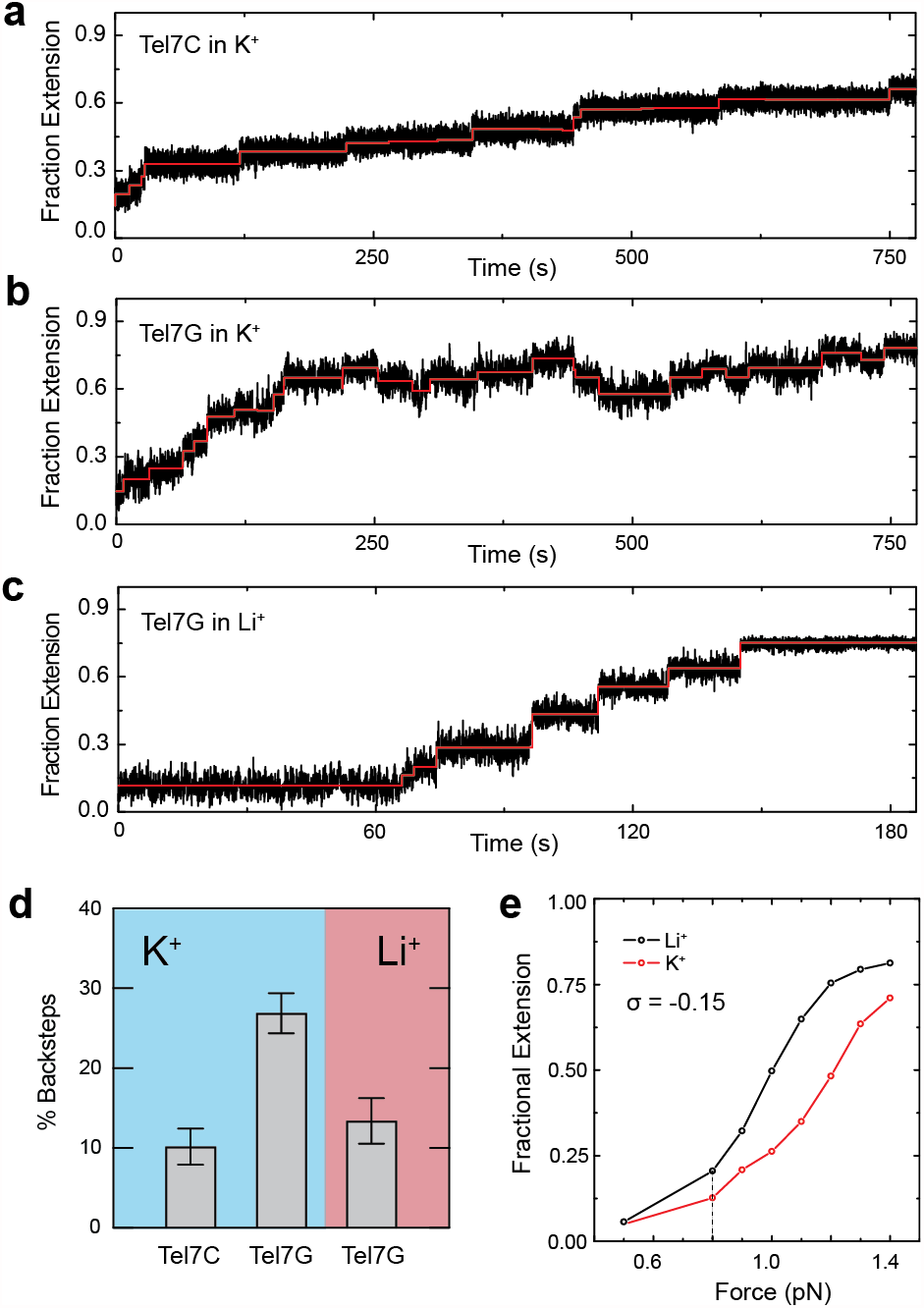
Sequence dependence of strand invasion time trace complexity. Representative telomere invasion time traces collected with Tel7C in K^+^ buffer (a), with Tel7G in K^+^ buffer (b), and with Tel7G in Li^+^ buffer (c). (d) Bar plot of the percent backstepping observed for Tel7C and Tel7G, in either K^+^ or Li^+^ buffer. Error bars are the standard deviation from each experiment repeated in triplicate, with the total number of steps for each replicate experiment, n ≥ 75. (e) Force extension curve of negatively supercoiled telomere DNA in Li^+^ and K^+^ held at *σ* = -0.15.

As a further test of this hypothesis, we next changed the buffer conditions to disfavor the stable folding of GQ structures by replacing K^+^ with Li^+^. We noted that the kinetics of Tel7G invasion in Li^+^ buffer was markedly faster compared to the identical experiment performed in K^+^ buffer (Figure 4A compared to Figure 4C). One possible explanation for this observation is that the target DNA duplex is less energetically stable in the presence of Li^+^ when compared to K^+^, a feature of Bform DNA that, to our knowledge, has not been biophysically characterized. To analyze this possibility, we compared the extension properties of the telomere DNA tethers as a function of superhelical density in both K^+^ and Li^+^. Indeed, we find that the DNA is more readily denatured by applied torques in the presence of Li^+^ (Figure 4E and Figure S7), which provides an explanation for the increased rate of tel7G invasion observed in our experiments. The effect of Li^+^ on the torsional stability of B-form DNA is not telomere-specific, as we observe a similar behavior for a non-telomeric DNA tether studied in K^+^ vs. Li^+^ (Figure S8). As described above, a prominent feature of the Tel7G invasion trajectories in K^+^ buffer is the prevalence of back-stepping throughout the invasion process. To quantify the frequency of backstepping within individual strand invasion experiments, we generate idealized trajectories using a step finding algorithm ^30^ and we define forward and backward steps as either an increase or decrease in extension in the DNA tether, respectively. Using this approach for the Tel7C strand, we find 10% ± 2% of the total steps to be backward steps, compared to 27% ± 3% of the total steps for the Tel7G strand (Figure 4A, 4B, and 4D). In contrast, performing the Tel7G strand invasion experiment in the presence of Li^+^ significantly reduced the frequency of backstepping to 13% ± 3%, a level comparable to that observed for the Tel7C experiments (Figure 4C and 4D). Taken together, these results again support a model wherein GQ folding in the displaced strand of a TDL underlies the complex dynamics and backstepping observed in Tel7G strand invasion trajectories in K^+^ conditions.

### Structural stability of telomeric D-loops formed by G-rich strand invasion

If the displaced strand within a TDL formed upon Grich strand invasion folds into a GQ structure, one prediction is that the TDL will be less energetically favored to resolve since the H-bonds that have been disrupted upon strand invasion are compensated by H-bonds within a GQ fold. As noted above, the stable unwinding of the telomere DNA target during strand invasion results in a change in the overall DNA twist (ie. the number of helical turns per unit length of the DNA molecule). Such changes in DNA twist, if structurally stable, can be directly measured as a shift in the rotation-extension curve to the left when the magnets are rotated back toward the relaxed state of the DNA molecule ^29^. In contrast, if the invading strands are ejected while rewinding the molecule back toward the relaxed state (ie. TDL resolution), one would expect to observe a rotation-extension curve that overlays the original pre-invaded state. After complete invasion of a target telomere DNA tether with the Tel7C oligonucleotide, the magnets were rotated back toward the relaxed state and into the positive superhelical density regime (Figure 5A). Overlay of multiple independent rotation-extension curves taken following Tel7C invasion reveal no detectable shift in the rotation extension curve (Figure 5A), consistent with the notion that as positive turns are being added back to the molecule the C-rich invasion strands are ejected from the molecule and the TDLs are resolved. In contrast, when the same experiment is conducted with the Tel7G invading strand, a significant shift in the rotation-extension curve is observed across many independent experimental trials (Figure 5B). This hysteresis in the rotation-extension curve taken on a molecule invaded by the Tel7G oligonucleotide is indicative of increased structural stability of TDLs when the physicologically relevant G-rich invading strand is used. Interestingly, if the same experiment is performed with the Tel7G strand in the presence of Li^+^ rather than K^+^, the observed hysteresis in the rotation-extension curve is eliminated (Figure 5C).

**Figure 5.**
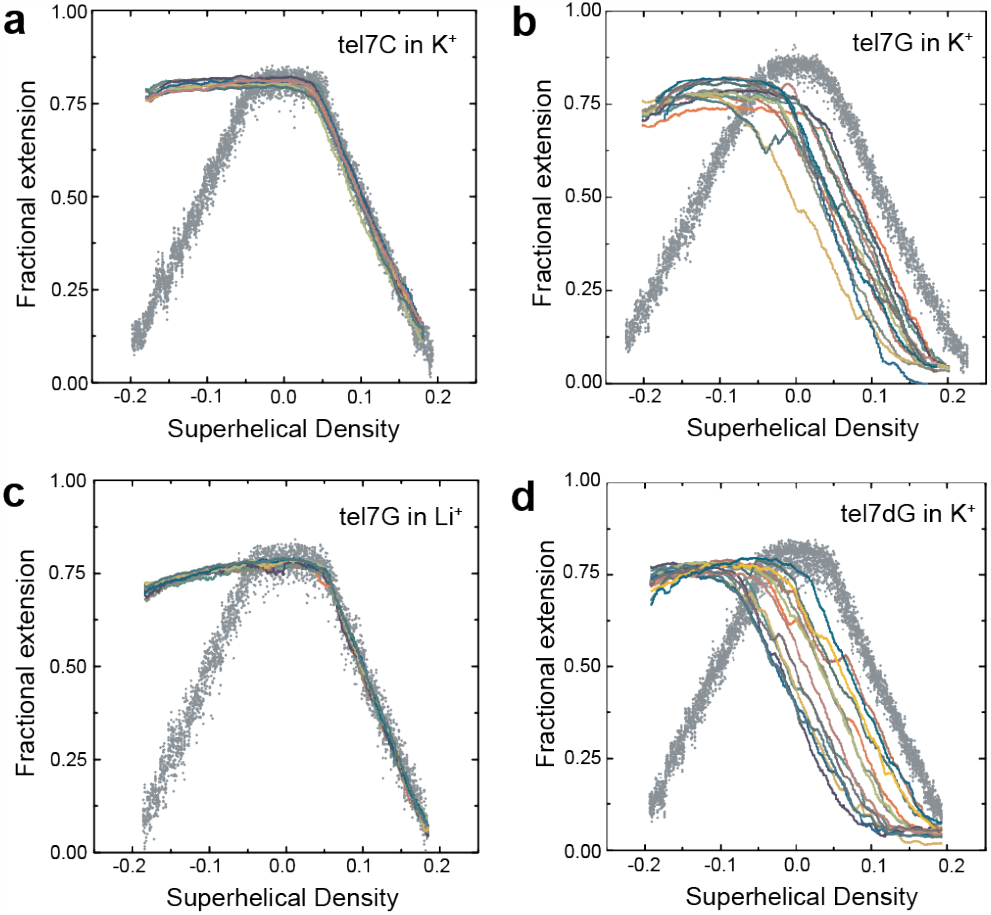
Stable telomeric D-loops result in a shift of the rotation extension curve. Initial rotation-extension curves in the absence of invading strands are shown in grey dots, while individual replicates (n=14) of rotation-extension data collected following complete strand invasion are shown in colored solid lines (n = 14 for all panels). (a) Tel7C in K^+^. (b) Tel7G in K^+^. (c) Tel7G in Li^+^. (d) Tel7dG in K^+^.

Although our results are consistent with a role for GQ in stabilizing TDL structures formed upon tel7G invasion, it remained a possiblity that the G-rich invading strands in solution were participating in intermolecular GQ formation with the diplaced strand, rather than formation of an intramolecular GQ within the displaced strand of the TDL. To distinguish between these two possibilities, we again turned to the use of a modified oligonucleotide wherein the central guanine in the G-run of every other telomere repeat was substituted by a 7-deazaguanine (Tel7dG), serving to disrupt the Hoogsteen face of the nucleoside. This modification prevents GQ folding while leaving the Watson-Crick face required for canonical helical basepairing unperturbed^33^. Analysis of DNA tethers following strand invastion by the Tel7dG strand again revealed a leftward shift of the rotation-extension curve, consistent with stable TDL formation (Figure 5D). These data lend further support to the notion that intramolecular GQs formed within the displaced strand of TDLs structurally stabilize the invaded state.

## DISCUSSION

Magnetic Tweezers (MT) force spectroscopy is a powerful tool with which to probe DNA mechanics ^34-36^. MT-based methods have been applied to the study of human telomere DNA in recent years, with a focus on the propensity of this repetitive G-rich sequence (TTAGGG)_n_ to fold into G-quadruplex (GQ) structures ^36-38^. Previously published single molecule spectroscopic analyses of telomere DNA mechanics have largely focused on the structural properties of short single-stranded (ss) model telomere DNA substrates ^36, 37, 39-44^. In the present work, we use a MT system to interrogate the structural properties of long, uninterrupted duplex telomere DNA molecules of physiologically relevant lengths (> 10 kilobases).

MT methods have also previously been used to directly monitor DNA strand invasion in real-time, providing a tool to study the mechanics of this essential DNA transaction that occurs during DNA repair and recombination pathways ^29, 45^. Here, we have adapted this approach to study strand invasion at telomere DNA target sites, a process proposed to occur during the formation of telomere-loops (T-loops) as well as during the alternative lengthening of telomeres (ALT) pathway ^15- 18^. By using telomeric ssDNA probes of various lengths introduced to individual duplex telomere DNA molecules held under precisely applied degrees of superhelical strain, we detect real-time strand invasion and the formation telomeric displacement-loops (TDLs). Varying the amount of applied torque to the target telomere DNA molecule reveals formation of TDLs is highly torque dependent. Analysis of the kinetics of strand invasion as a function of applied torque using a simple transition state model ^32^ suggests ∼9 base pairs of the DNA target must be unwound at the transition state along the well-defined strand invasion reaction coordinate. This result supports a previous model for the role of the telomere repeat binding factor 2 protein (TRF2), which has been shown to wrap duplex telomere DNA in a chiral fashion, resulting in the application of negative superhelical strain and promoting T-loop formation ^46, 47^.

Interestingly, we observe complex invasion dynamics when the invasion trajectories are collected in the presence of a G-rich ssDNA oligonucleotide, intended to model the G-rich 3’ ssDNA tail that exists at endogenous telomere ends. The invasion dynamics are characterized by a combination of forward and reverse steps, and these back steps are suppressed when performing the same experiments with the complementary C-rich strand or in the presence of Li^+^ rather than K^+^ ions. It is well established that Li^+^ has a destabilizing effect on GQ folding ^21^. We also provide evidence that TDLs formed upon G-rich strand invasion in the presence of K^+^ are more energetically stable than when formed in the presence of Li^+^ or with the C-rich strand. Taken together, these results lead to a model wherein the formation of a TDL upon invasion of the G-rich ssDNA tail permits the G-rich displaced strand to fold into a GQ structure (Figure 6, Figure S9). While it is well documented that single-stranded telomere DNA substrates fold into GQ structures *in vitro* ^21, 48-51^, the prevalence of this structure at telomeres and elsewhere within the genome has been the subject of debate ^23, 52^. Interestingly, a recently reported magnetic tweezers study of the promoter region of the c-kit oncogene demonstrated that negative superhelical strain can also drive the B-form to GQ structural transition ^53^. The results of our mechanical analysis of telomeric D-loops suggest the process of strand invasion at telomere DNA targets may provide an opportunity for GQ structures to fold *in vivo*, as has recently been reported by live cell imaging ^54, 55^.

**Figure 6.**
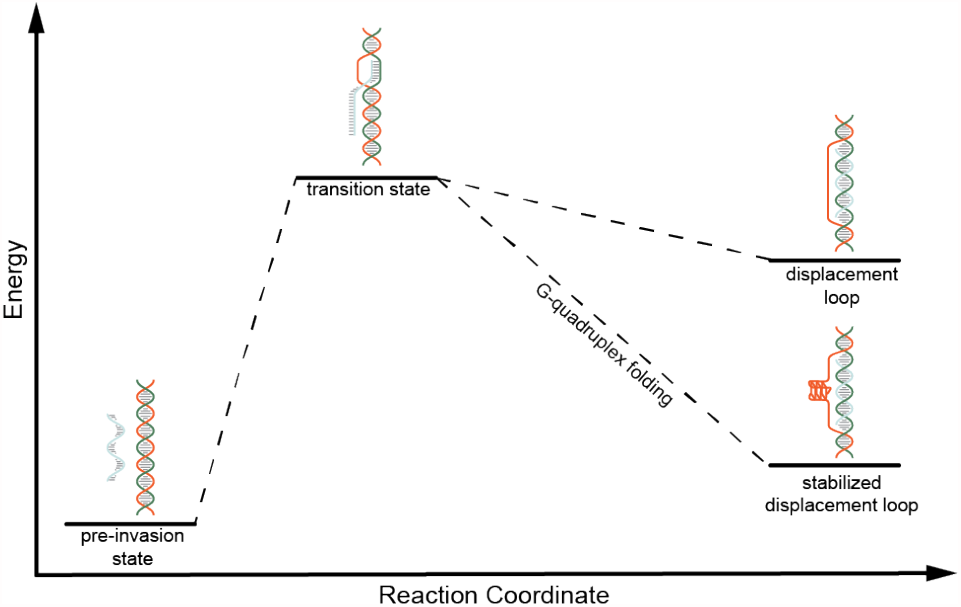
Model of telomeric displacement loop stabilization by G-quadruplex folding within the displaced strand. Under GQ forming conditions, the displaced G-rich strand of the TDL can fold into a GQ stabilizing the overall structure.

## CONCLUSIONS

The system we describe in the present study provides a powerful experimental platform for future studies of strand invasion at telomere DNA targets. For example, single-molecule studies using this system can be designed to understand the molecular mechanisms of telomere-associate proteins and enzymes known to resolve D-loop and GQ structures ^56-58^. Moreover, our novel system can be employed to directly study the mechanism of GQ-binding compounds and their possible role in stabilizing TDLs^59^. Lastly, recent studies have shown that telomeres, long thought to be transcriptionally silent, are transcribed to generate long non-coding Telomere repeat-containing RNA (TERRA)^60^. TERRA is implicated in regulating various aspects of telomere biology and is proposed to do so through the formation of RNA-loops (R-loops) at telomeres ^61^. Thus, future work utilizing our novel MT-based assay will also focus on the mechanical properties of telomeric R-loops and the molecular mechanism of TERRA-mediated regulation of telomere function.

## ASSOCIATED CONTENT

### Supporting Information

Detailed materials and methods and figures S1-S9 can be found in the supplemental information document.

## AUTHOR INFORMATION

### Funding Sources

This work was supported by NIH grant R01-GM095850 to M.D.S and National Science Foundation GRFP (DGE 0809125) and Eugene Cota-Robles Fellowship to J.W.P.

## ACKNOWLEDGMENT

We thank Professor Clive Bagshaw for helpful conversations during preparation of this manuscript. We thank Professor Jack Griffith (UNC Chapel Hill) for the gift of the pRST5 plasmid.

## Notes

### Competing Interest Statement

The authors have declared no competing interest.

## REFERENCES

1. Palm, W.; de Lange, T.; How shelterin protects mammalian telomeres. Annu Rev Genet 2008, 42, 301–34.

2. de Lange, T.; Shelterin-Mediated Telomere Protection. Annu Rev Genet 2018, 52, 223-247.

3. Hastie, N. D.; Dempster, M.; Dunlop, M. G.; Thompson, A. M.; Green, D. K.; Allshire, R. C.; Telomere reduction in human colorectal carcinoma and with ageing. Nature 1990, 346 (6287), 866–8.

4. Lansdorp, P. M.; Verwoerd, N. P.; van de Rijke, F. M.; Dragowska, V.; Little, M. T.; Dirks, R. W.; Raap, A. K.; Tanke, H. J.; Heterogeneity in telomere length of human chromosomes. Hum Mol Genet 1996, 5 (5), 685–91.

5. Makarov, V. L.; Hirose, Y.; Langmore, J. P.; Long G tails at both ends of human chromosomes suggest a C strand degradation mechanism for telomere shortening. Cell 1997, 88 (5), 657–66.

6. Olovnikov, A. M.; A theory of marginotomy. The incomplete copying of template margin in enzymic synthesis of polynucleotides and biological significance of the phenomenon. J Theor Biol 1973, 41 (1), 181–90.

7. Prelich, G.; Stillman, B.; Coordinated leading and lagging strand synthesis during SV40 DNA replication in vitro requires PCNA. Cell 1988, 53 (1), 117–26.

8. Hayflick, L.; Moorhead, P. S.; The serial cultivation of human diploid cell strains. Exp Cell Res 1961, 25, 585–621.

9. Kim, N. W.; Piatyszek, M. A.; Prowse, K. R.; Harley, C. B.; West, M. D.; Ho, P. L.; Coviello, G. M.; Wright, W. E.; Weinrich, S. L.; Shay, J. W.; Specific association of human telomerase activity with immortal cells and cancer. Science 1994, 266 (5193), 2011–5.

10. Greider, C. W.; Blackburn, E. H.; Identification of a specific telomere terminal transferase activity in Tetrahymena extracts. Cell 1985, 43 (2 Pt 1), 405–13.

11. Greider, C. W.; Blackburn, E. H.; The telomere terminal transferase of Tetrahymena is a ribonucleoprotein enzyme with two kinds of primer specificity. Cell 1987, 51 (6), 887-98.

12. Greider, C. W.; Blackburn, E. H.; A telomeric sequence in the RNA of Tetrahymena telomerase required for telomere repeat synthesis. Nature 1989, 337 (6205), 331–7.

13. Weinrich, S. L.; Pruzan, R.; Ma, L.; Ouellette, M.; Tesmer, V. M.; Holt, S. E.; Bodnar, A. G.; Lichtsteiner, S.; Kim, N. W.; Trager, J. B.; Taylor, R. D.; Carlos, R.; Andrews, W. H.; Wright, W. E.; Shay, J. W.; Harley, C. B.; Morin, G. B.; Reconstitution of human telomerase with the template RNA component hTR and the catalytic protein subunit hTRT. Nat Genet 1997, 17 (4), 498-502.

14. Dilley, R. L.; Greenberg, R. A.; ALTernative Telomere Maintenance and Cancer. Trends Cancer 2015, 1 (2), 145–156.

15. Zhang, J. M.; Yadav, T.; Ouyang, J.; Lan, L.; Zou, L.; Alternative Lengthening of Telomeres through Two Distinct Break-Induced Replication Pathways. Cell Rep 2019, 26 (4), 955–968 e3.

16. Griffith, J. D.; Comeau, L.; Rosenfield, S.; Stansel, R. M.; Bianchi, A.; Moss, H.; de Lange, T.; Mammalian telomeres end in a large duplex loop. Cell 1999, 97 (4), 503–14.

17. Tomaska, L.; Nosek, J.; Kar, A.; Willcox, S.; Griffith, J. D.; A New View of the T-Loop Junction: Implications for Self-Primed Telomere Extension, Expansion of Disease-Related Nucleotide Repeat Blocks, and Telomere Evolution. Front Genet 2019, 10, 792.

18. Greider, C. W.; Telomeres do D-loop-T-loop. Cell 1999, 97 (4), 419–22.

19. Dilley, R. L.; Verma, P.; Cho, N. W.; Winters, H. D.; Wondisford, A. R.; Greenberg, R. A.; Break-induced telomere synthesis underlies alternative telomere maintenance. Nature 2016, 539 (7627), 54–58.

20. Zhao, Y.; Sfeir, A. J.; Zou, Y.; Buseman, C. M.; Chow, T. T.; Shay, J. W.; Wright, W. E.; Telomere extension occurs at most chromosome ends and is uncoupled from fillin in human cancer cells. Cell 2009, 138 (3), 463–75.

21. Lane, A. N.; Chaires, J. B.; Gray, R. D.; Trent, J. O.; Stability and kinetics of Gquadruplex structures. Nucleic Acids Res 2008, 36 (17), 5482–515.

22. Lee, J. Y.; Yoon, J.; Kihm, H. W.; Kim, D. S.; Structural diversity and extreme stability of unimolecular Oxytricha nova telomeric Gquadruplex. Biochemistry 2008, 47 (11), 3389–96.

23. Huppert, J. L.; Balasubramanian, S.; Prevalence of quadruplexes in the human genome. Nucleic Acids Res 2005, 33 (9), 2908–16.

24. Balasubramanian, S.; Neidle, S.; Gquadruplex nucleic acids as therapeutic targets. Curr Opin Chem Biol 2009, 13 (3), 345–53.

25. Stansel, R. M.; de Lange, T.; Griffith, J. D.; Tloop assembly in vitro involves binding of TRF2 near the 3’ telomeric overhang. EMBO J 2001, 20 (19), 5532–40.

26. Bustamante, C.; Marko, J. F.; Siggia, E. D.; Smith, S.; Entropic elasticity of lambda-phage DNA. Science 1994, 265 (5178), 1599–600.

27. Baumann, C. G.; Smith, S. B.; Bloomfield, V. A.; Bustamante, C.; Ionic effects on the elasticity of single DNA molecules. Proc Natl Acad Sci U S A 1997, 94 (12), 6185–90.

28. Buglione, E.; Salerno, D.; Marrano, C. A.; Cassina, V.; Vesco, G.; Nardo, L.; Dacasto, M.; Rigo, R.; Sissi, C.; Mantegazza, F.; Nanomechanics of G-quadruplexes within the promoter of the KIT oncogene. Nucleic Acids Res 2021, 49 (8), 4564–4573.

29. Strick, T. R.; Croquette, V.; Bensimon, D.; Homologous pairing in stretched supercoiled DNA. Proc Natl Acad Sci U S A 1998, 95 (18), 10579–83.

30. Kerssemakers, J. W.; Munteanu, E. L.; Laan, L.; Noetzel, T. L.; Janson, M. E.; Dogterom, M.; Assembly dynamics of microtubules at molecular resolution. Nature 2006, 442 (7103), 709–12.

31. Strick, T.; Allemand, J.; Croquette, V.; Bensimon, D.; Twisting and stretching single DNA molecules. Prog Biophys Mol Biol 2000, 74 (1-2), 115–40.

32. Ramreddy, T.; Sachidanandam, R.; Strick, T. R.; Real-time detection of cruciform extrusion by single-molecule DNA nanomanipulation. Nucleic Acids Res 2011, 39 (10), 4275–83.

33. Masai, H.; Kakusho, N.; Fukatsu, R.; Ma, Y.; Iida, K.; Kanoh, Y.; Nagasawa, K.; Molecular architecture of G-quadruplex structures generated on duplex Rif1-binding sequences. J Biol Chem 2018, 293 (44), 17033–17049.

34. Strick, T. R.; Allemand, J. F.; Bensimon, D.; Croquette, V.; Behavior of supercoiled DNA. Biophys J 1998, 74 (4), 2016–28.

35. Yu, Z.; Dulin, D.; Cnossen, J.; Kober, M.; van Oene, M. M.; Ordu, O.; Berghuis, B. A.; Hensgens, T.; Lipfert, J.; Dekker, N. H.; A force calibration standard for magnetic tweezers. Rev Sci Instrum 2014, 85 (12), 123114.

36. Long, X.; Parks, J. W.; Stone, M. D.; Integrated magnetic tweezers and singlemolecule FRET for investigating the mechanical properties of nucleic acid. Methods 2016, 105, 16–25.

37. You, H.; Le, S.; Chen, H.; Qin, L.; Yan, J.; Single-molecule Manipulation of Gquadruplexes by Magnetic Tweezers. J Vis Exp 2017, (127).

38. Lin, J.; Kaur, P.; Countryman, P.; Opresko, P. L.; Wang, H.; Unraveling secrets of telomeres: one molecule at a time. DNA Repair (Amst) 2014, 20, 142–153.

39. An, N.; Fleming, A. M.; Middleton, E. G.; Burrows, C. J.; Single-molecule investigation of G-quadruplex folds of the human telomere sequence in a protein nanocavity. Proc Natl Acad Sci U S A 2014, 111 (40), 14325–31.

40. Hwang, H.; Kreig, A.; Calvert, J.; Lormand, J.; Kwon, Y.; Daley, J. M.; Sung, P.; Opresko, P. L.; Myong, S.; Telomeric overhang length determines structural dynamics and accessibility to telomerase and ALT-associated proteins. Structure 2014, 22 (6), 842–53.

41. Lee, J. Y.; Okumus, B.; Kim, D. S.; Ha, T.; Extreme conformational diversity in human telomeric DNA. Proc Natl Acad Sci U S A 2005, 102 (52), 18938–43.

42. Long, X.; Parks, J. W.; Bagshaw, C. R.; Stone, M. D.; Mechanical unfolding of human telomere G-quadruplex DNA probed by integrated fluorescence and magnetic tweezers spectroscopy. Nucleic Acids Res 2013, 41 (4), 2746–55.

43. Parks, J. W.; Stone, M. D.; Single-Molecule Studies of Telomeres and Telomerase. Annu Rev Biophys 2017, 46, 357–377.

44. Ray, S.; Bandaria, J. N.; Qureshi, M. H.; Yildiz, A.; Balci, H.; G-quadruplex formation in telomeres enhances POT1/TPP1 protection against RPA binding. Proc Natl Acad Sci U S A 2014, 111 (8), 2990–5.

45. Lee, M.; Lipfert, J.; Sanchez, H.; Wyman, C.; Dekker, N. H.; Structural and torsional properties of the RAD51-dsDNA nucleoprotein filament. Nucleic Acids Res 2013, 41 (14), 7023–30.

46. Amiard, S.; Doudeau, M.; Pinte, S.; Poulet, A.; Lenain, C.; Faivre-Moskalenko, C.; Angelov, D.; Hug, N.; Vindigni, A.; Bouvet, P.; Paoletti, J.; Gilson, E.; Giraud-Panis, M. J.; A topological mechanism for TRF2enhanced strand invasion. Nat Struct Mol Biol 2007, 14 (2), 147–54.

47. Benarroch-Popivker, D.; Pisano, S.; Mendez-Bermudez, A.; Lototska, L.; Kaur, P.; Bauwens, S.; Djerbi, N.; Latrick, C. M.; Fraisier, V.; Pei, B.; Gay, A.; Jaune, E.; Foucher, K.; Cherfils-Vicini, J.; Aeby, E.; Miron, S.; Londono-Vallejo, A.; Ye, J.; Le Du, M. H.; Wang, H.; Gilson, E.; Giraud-Panis, M. J.; TRF2-Mediated Control of Telomere DNA Topology as a Mechanism for Chromosome-End Protection. Mol Cell 2016, 61 (2), 274–86.

48. Ambrus, A.; Chen, D.; Dai, J.; Bialis, T.; Jones, R. A.; Yang, D.; Human telomeric sequence forms a hybrid-type intramolecular G-quadruplex structure with mixed parallel/antiparallel strands in potassium solution. Nucleic Acids Res 2006, 34 (9), 2723–35.

49. Li, J.; Correia, J. J.; Wang, L.; Trent, J. O.; Chaires, J. B.; Not so crystal clear: the structure of the human telomere G-quadruplex in solution differs from that present in a crystal. Nucleic Acids Res 2005, 33 (14), 4649–59.

50. Parkinson, G. N.; Lee, M. P.; Neidle, S.; Crystal structure of parallel quadruplexes from human telomeric DNA. Nature 2002, 417 (6891), 876–80.

51. Wang, Y.; Patel, D. J.; Solution structure of the human telomeric repeat d[AG3(T2AG3)3] Gtetraplex. Structure 1993, 1 (4), 263–82.

52. Rhodes, D.; Lipps, H. J.; G-quadruplexes and their regulatory roles in biology. Nucleic Acids Res 2015, 43 (18), 8627–37.

53. Buglione, E.; Salerno, D.; Marrano, C. A.; Cassina, V.; Vesco, G.; Nardo, L.; Dacasto, M.; Rigo, R.; Sissi, C.; Mantegazza, F.; Nanomechanics of G-quadruplexes within the promoter of the KIT oncogene. Nucleic Acids Res 2021.

54. Di Antonio, M.; Ponjavic, A.; Radzevicius, A.; Ranasinghe, R. T.; Catalano, M.; Zhang, X.; Shen, J.; Needham, L. M.; Lee, S. F.; Klenerman, D.; Balasubramanian, S.; Singlemolecule visualization of DNA G-quadruplex formation in live cells. Nat Chem 2020, 12 (9), 832–837.

55. Summers, P. A.; Lewis, B. W.; Gonzalez-Garcia, J.; Porreca, R. M.; Lim, A. H. M.; Cadinu, P.; Martin-Pintado, N.; Mann, D. J.; Edel, J. B.; Vannier, J. B.; Kuimova, M. K.; Vilar, R.; Visualising G-quadruplex DNA dynamics in live cells by fluorescence lifetime imaging microscopy. Nat Commun 2021, 12 (1), 162.

56. Wu, Y.; Shin-ya, K.; Brosh, R. M., Jr.; FANCJ helicase defective in Fanconia anemia and breast cancer unwinds G-quadruplex DNA to defend genomic stability. Mol Cell Biol 2008, 28 (12), 4116–28.

57. Budhathoki, J. B.; Maleki, P.; Roy, W. A.; Janscak, P.; Yodh, J. G.; Balci, H.; A Comparative Study of G-Quadruplex Unfolding and DNA Reeling Activities of Human RECQ5 Helicase. Biophys J 2016, 110 (12), 2585–2596.

58. Tippana, R.; Hwang, H.; Opresko, P. L.; Bohr, V. A.; Myong, S.; Single-molecule imaging reveals a common mechanism shared by G-quadruplex-resolving helicases. Proc Natl Acad Sci U S A 2016, 113 (30), 8448–53.

59. Tan, J.; Lan, L.; The DNA secondary structures at telomeres and genome instability. Cell Biosci 2020, 10, 47.

60. Azzalin, C. M.; Reichenbach, P.; Khoriauli, L.; Giulotto, E.; Lingner, J.; Telomeric repeat containing RNA and RNA surveillance factors at mammalian chromosome ends. Science 2007, 318 (5851), 798–801.

61. Oliva-Rico, D.; Herrera, L. A.; Regulated expression of the lncRNA TERRA and its impact on telomere biology. Mech Ageing Dev 2017, 167, 16–23.

