## Supplementary Information for "Single-molecule Mechanical Analysis of Strand Invasion in Human Telomere DNA"

Detailed Experimental Methods

Table S1. Oligonoucleotides Table.

Figure S1. Rotation Extension Curves of Telomere DNA and Lambda DNA in $K^+$

Figure S2. Negatively supercoiled telomeric DNA in the absence of invading oligonucleotides.

Figure S3. Representative Tel7C strand invasion traces.

Figure S4. Tel7C step size histograms using different length oligonucleotides for invasion.

Figure S5. Dwell time distribution analysis for Tel7C and Tel7G strand invasion traces.

Figure S6. Representative Tel7dG strand invasion traces.

Figure S7. Telomere DNA torque response in $K^+$ and $Li^+$ at various forces.

Figure S8. Lambda phage DNA torque response in $K^+$ and $Li^+$ at various forces.

Figure S9. Model of a telomeric D-loop (TDL).

### MATERIALS AND METHODS

#### DNA oligonucleotides

Custom synthetic DNA oligonucleotides were purchased from Integrated DNA Technologies, Inc or for the 7deazaG oligonucleotide from Midland Certified Reagent Company. The sequences of all DNAs used in the study are listed in Table S1.

#### DNA molecules for magnetic tweezers

The telomere molecules consist of double stranded repeat telomere DNA sequence (~7000bp – 13000bp) flanked by two DNA linkers. The linkers are synthesized by PCR using Taq polymerase (New England Biolabs) with a pUC19 template using primers flanking the multiple cloning site (see Supplementary Table 1). Each PCR was set up using a dNTP mixture containing a 1:9 molar ratio of digoxygenin-11-dUTP (Roche) or biotin-11-dUTP (Roche) to dTTP followed by digestion with BamH1 to generate a ~500bp linker. To these linkers, DNA adapters constructed by annealing two oligonucleotides (see Supplementary Table 1) are ligated to the BamH1 overhang that generate overhangs complimentary to the telomere fragments, followed by gel purification (Qiagen). The telomere fragments are generated by sequential digestion of the pRST5 plasmid with Bbs1 and Bsmb1 as described [1], generating a ~576bp double stranded telomeric DNA fragment with a 5' AGGG overhang and a 3' TCCC overhang, followed by gel purification (Qiagen).

The molecules are assembled by first ligating ~1pmole of the digoxygenin-labeled linker to the telomere fragment in a 1:5 molar ratio at 16°C for 1 hour. After an hour, ~16pmoles of the 576bp telomere fragment are added to the ligation reaction in 1X T4 DNA ligase buffer with T4 ligase and incubated at 16°C for ~4 hours. After 4 hours, ~3pmoles of the biotin-labeled linker are added to the reaction in 1X T4 DNA ligase buffer with T4 ligase and incubated at 16°C overnight. The molecules are then phenol/chloroform extracted and ethanol precipitated to reduce the volume to facilitate gel extraction. The molecules are then gel purified (Qiagen) away from unwanted circular byproducts on a 0.6 % agarose gel stained with SYBR® safe DNA stain illuminated under blue light to avoid UV light exposure which could damage the DNA. The section of gel excised corresponded to DNA lengths between 6kb and the well. Following gel extraction, the molecules are preCR® (New England Biolabs) treated to fill any nicks or gaps that could have formed during the construction, followed by a subsequent phenol/chloroform extraction and ethanol precipitation.

The lambda phage DNA molecules consist of ~11kbp of a section of lambda phage DNA that was selected to have 50% GC content and is free from BamH1 and HindIII restriction enzyme cut sites. The fragment is first amplified by PCR using full length lambda phage DNA as a template (NEB). The primers used for PCR each contain either a BamH1 or HindIII restriction enzyme cut site. The PCR product is then phenol chloroform extracted then ethanol precipitated and resuspended in water. The resuspended DNA is subject to restriction enzyme digestion using BamH1 and HindIII, followed by another round of phenol – chloroform extraction then ethanol precipitation and resuspended in water. The lambda phage DNA fragment is then ligated to similar biotin- or digoxygenin-labeled linker DNA material used as described above for the telomere molecule construction, only these linkers contain either BamH1 or HindIII complimentary sticky ends. The ligation product is then gel purified to remove unwanted linker DNA material.

### Flow cell assembly and functionalization

A glass coverslip (Fischer Scientific, Cat. No. 12-548-5E) is first washed with a 5% Alconox® solution, extensively rinsed with $ddH_2O$ and dried under nitrogen. It is then plasma cleaned (Harrick, PDC-3XG) for ~5 minutes with air plasma. Once cleaned, 200µL of a 0.2% nitrocellulose (Biorad Cat. No. 162-0115) solution dissolved in absolute ethanol is applied by spin coating for 30 seconds at 6k rpm and cured at 180°C for 1.5 minutes on a hot plate. Flow cells are assembled by cutting a ~20µL channel in a piece of double-sided tape sandwiched between the nitrocellulose coated coverslip and another cleaned coverslip with two holes drilled at each end of the channel. A 100µL reservoir made from a cut pipette tip is glued at one hole and a piece of tubing (Intramedic® Cat. No. 427426) at the other to act as a fluidics system for buffer exchange.

### Immobilization of DNA molecules

To the flow cell, 20µL of 200mg/mL anti-digoxygenin antibody (Roche, Cat. No. 11333089001) is incubated at 4°C overnight. The next morning, 1µm polystyrene reference beads (Spherotech Cat. # AP10-10) diluted 1:1000 in PBS pH 7.4 is added and allowed to incubate for 8 minutes. The channel is then blocked for 1 hour using 100µL of 5mg/mL BSA in PBS pH 7.4. Telomeric DNA constructs diluted 1:50 into the same blocking buffer is then introduced to the flow cell and allowed to incubate for 1 hour. 1µm streptavidin coated magnetic beads (Invitrogen, Cat. No. 65601) pre-blocked and diluted 1:500 in blocking buffer is then introduced to the flow cell and allowed to incubate for 1 hour. Finally, ~1mL imaging buffer (150mM $K^+$ or $Li^+$, 10mM TRIS, 0.1mg/mL yeast tRNA) is flowed through the flow cell to clear the channel of free DNA and beads.

### Magnetic tweezers force calibration

The magnetic tweezers microscope used in this study is previously described [2, 3], programs used for data collection and analysis were written in house using LabVIEW and Matlab and are available upon request.

To calibrate the forces applied to each DNA tether, forces were measured across a wide range of magnet positions above the flow cell. Data is collected with an acquisition time of 490Hz. Forces applied to the tether were determined by tracking the variance of the bead position in the x-axis ($\langle \chi^2 \rangle$) and calculated using the expression $F = Lk_bT/\langle \chi^2 \rangle$, where $L$ is the contour length of the DNA molecule and $k_bT$ is thermal energy (4.1pN × nm) at room temperature. The DNA tether length was measured by calibrating the diffraction ring pattern of the magnetic bead using a piezo-controlled objective positioning device (Mad City Labs) as previously described[4]. The force vs. magnet position was plotted and fit to a double exponential decay function to generate of force calibration as described[4]. Force calibrations were performed individually for every DNA tether used in this study. Force vs. extension curves were then generated and fit to the worm-like chain polymer model as described [5], allowing for the determination of the persistence length for a given molecule, which acts as a DNA quality check.

### Experimental procedure

Once a molecule is selected for study, an extension vs. superhelical density curve is generated at the desired force set point determined for the experiment. Data is collected with an acquisition time of 100Hz. The negative superhelical density region of the curve is visually inspected for linearity, and the lowest superhelical density still corresponding to the linear region is the amount of turns applied to the molecule before inducing strand invasion. Once determined, the molecule is returned to a relaxed state and held at ~5pN while >10 chamber volumes of oligonucleotides in either $K^+$ or $Li^+$ imaging buffer is introduced into the flow cell. The molecule is then held at ~0.3pN, supercoiled to the predetermined superhelical density, then pulled at the desired force set point. We set time equal to zero once the magnets reach the position corresponding to the force set point. We allow strand invasion to occur until the extension of the molecule reaches ~85% of the extension at the force set point, at which point the molecule is overwound to double the amount of turns originally imparted on the molecule and the extension vs. superhelical density is recorded. Once the overwind is complete, the molecule is pulled at maximum force (which varies for each tether but is typically between 15 – 25pN) to eject any invaded oligonucleotides so the experiment can be repeated.

**Step fitting**

Steps were fit to invasion time traces using a previously described algorithm [6]. Data was intentionally slightly overfit to ensure no steps were missed. The tel7C data was used to calculate the standard deviation of the step size for that length invading DNA oligonucleotide, and steps that were found to be smaller than two times the standard deviation were removed from the data. All Matlab scripts used were written in house and available upon request.

### Table S1

| Oligo Name | Sequence (5' - 3') |
|---|---|
| PCR Primer 1 for Biotin/Digoxygenin labeled DNA linker | ACATTTCCCCGAAAAGTGCCA |
| PCR Primer 2 for Biotin/Digoxygenin labeled DNA linker | TCAAGTCAGAGGTGGCGAAAC |
| BamH1 overhang for adapter ligation | /5Phos/GATCCAGTCTGCGTACAGTGG |
| CCCT telomere overhang for adapter ligation | /5Phos/CCCTCCACTGTACGCAGACTG ' |
| AGGG telomere overhang for adapter ligation | /5Phos/AGGGCCACTGTACGCAGACTG |
| Tel3C | AATCCCAATCCCAATCCCAATCCCAATCCCAATCCCAATCCC |
| Tel7C | AATCCCAATCCCAATCCCAATCCCAATCCCAATCCCAATCCC |
| Tel715C | AATCCCAATCCCAATCCCAATCCCAATCCCAATCCCAATCCCAATCCCA ATCCCAATCCCAATCCCAATCCCAATCCCAATCCCAATCCC |
| Tel7G | TTAGGGTTAGGGTTAGGGTTAGGGTTAGGGTTAGGGTTAGGG |
| Tel7dG | TTAG(7-deaza-dG)GTTAGGGTTAG(7-deaza-dG)GTTAGGGTTAG(7-deaza-dG)GTTAGGGTTAG(7-deaza-dG)G |
| PCR primer 1 for lambda phage DNA molecule | GATCGGATCCGTAAGGGATGTTTATGACGAGC |
| PCR primer 2 for lambda phage DNA molecule | GATCAAGCTTTCAGCTCCCATACGCTGTATTC |

**Figure S1**

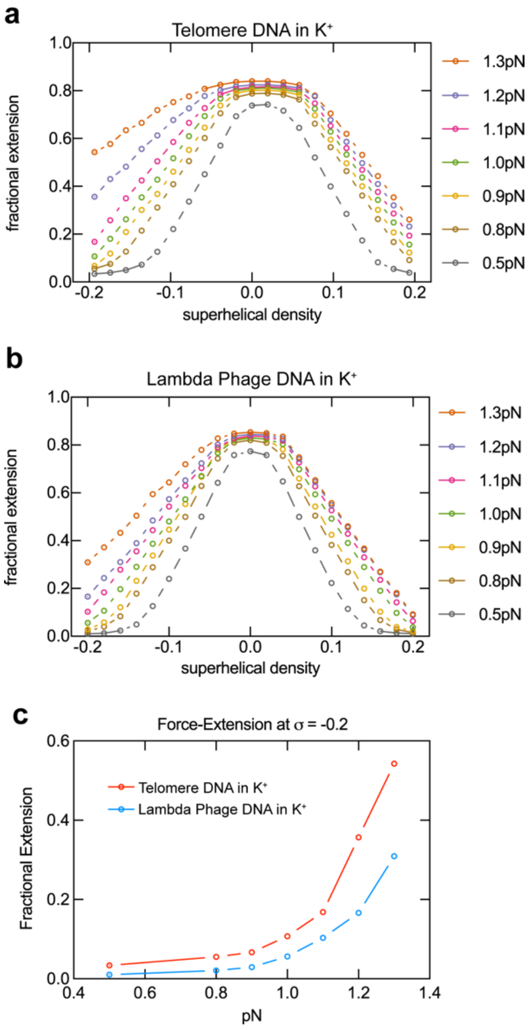


Figure S1. Rotation Extension Curves of telomere DNA (a) and lambda phage DNA (b) in $K^+$. C) Force-extension curve of telomere DNA (red) and lambda phage DNA (blue) at a superhelical density of -0.2.

**Figure S2**

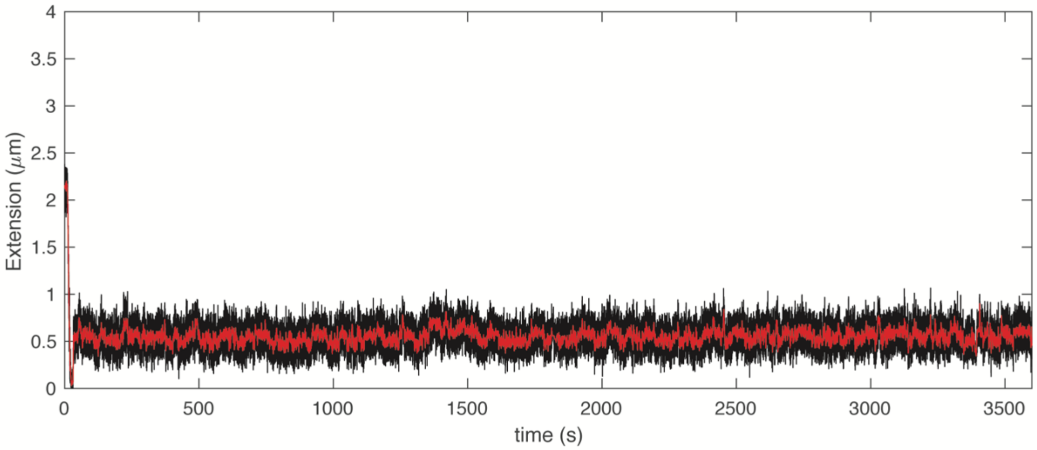


Figure S2. Negatively supercoiled telomeric DNA in the absence of invading strand shows no change in extension signal. Change in extension at the beginning of the trace corresponds to introduction of negative supercoils by rotating the magnets at the start of the experiment.

**Figure S3**

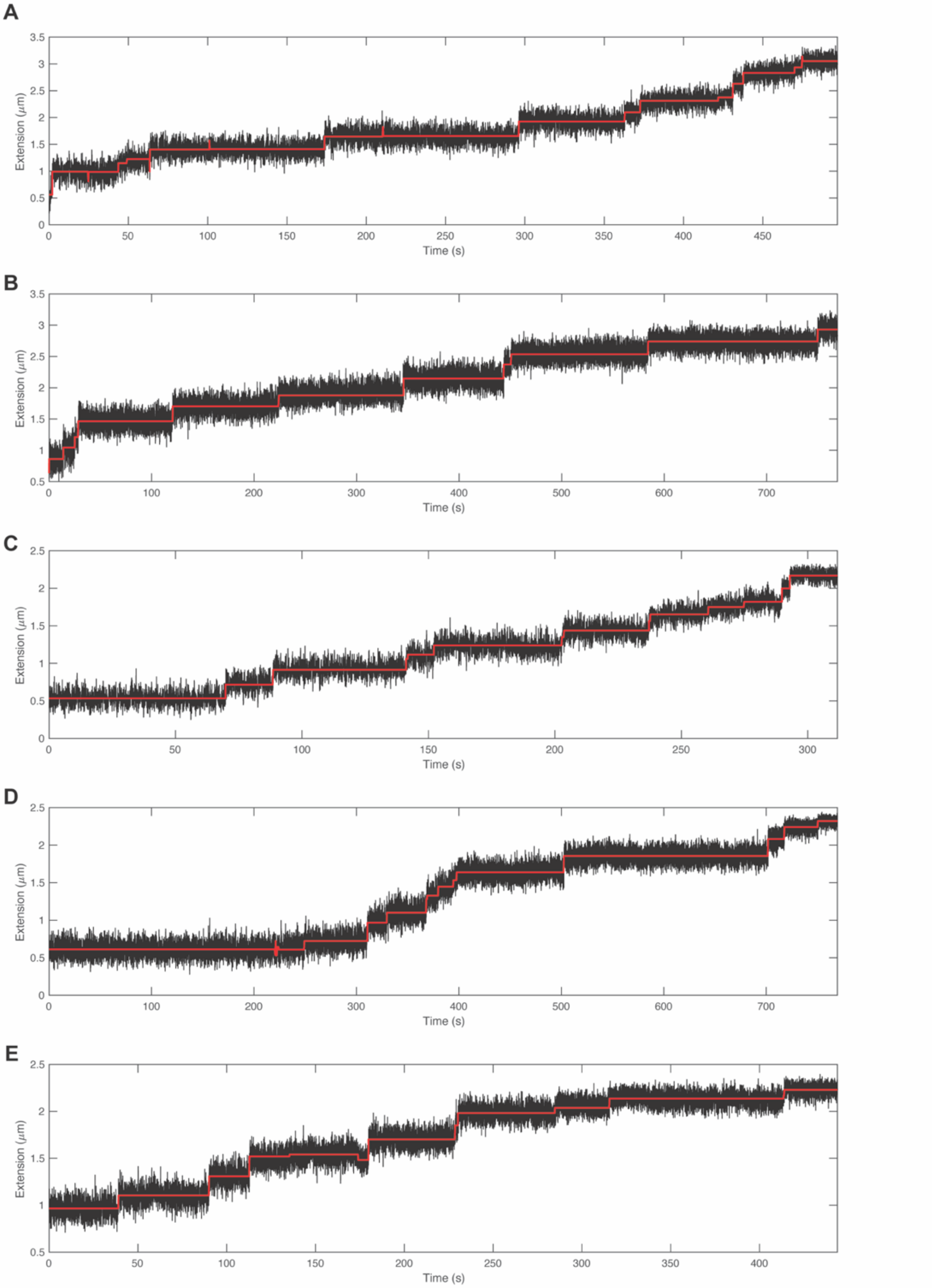


Figure S3. Representative Tel7C strand invasion traces. Raw data collected at 100 Hz are shown in black and idealized trace output from a stepfitting algorithm shown in red [6].

### Figure S4

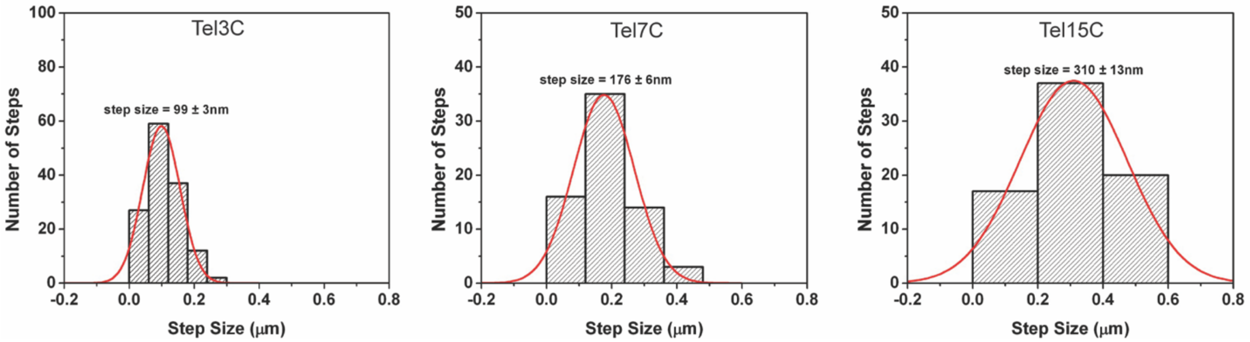


Figure S4. Tel7C step size histograms using different length oligonucleotides for invasion. Gaussian fits shown in red to calculate the mean step size.

**Figure S5**

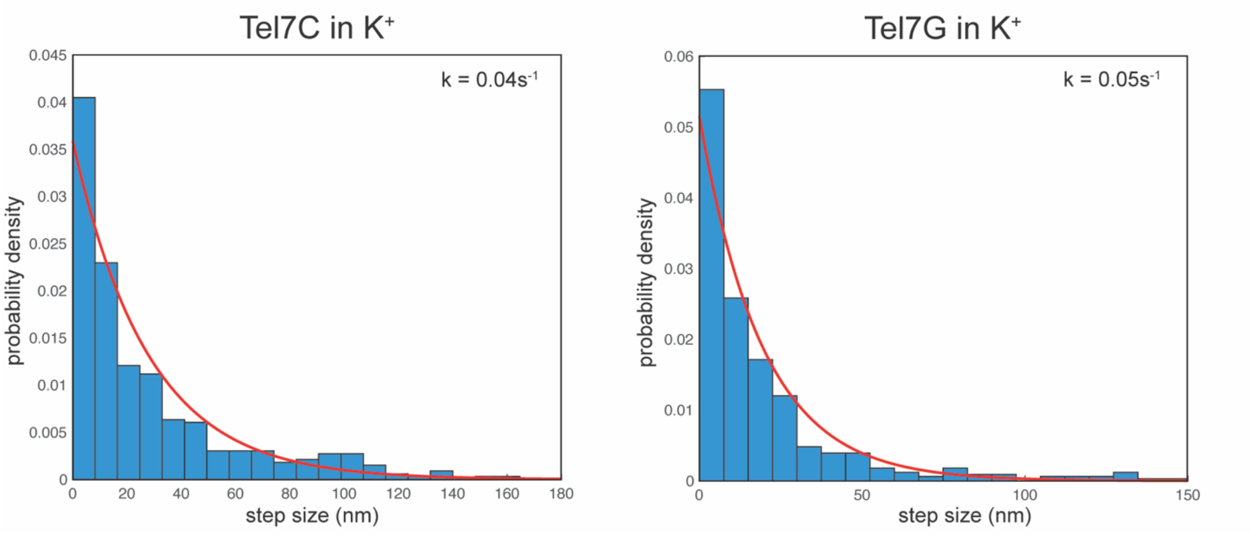


Figure S5. Dwell time distribution analysis for Tel7C and Tel7G strand invasion traces. Each data set was fit to a single exponential decay function using MEMLET [7] (red line) to calculate the rate constant.

**Figure S6**

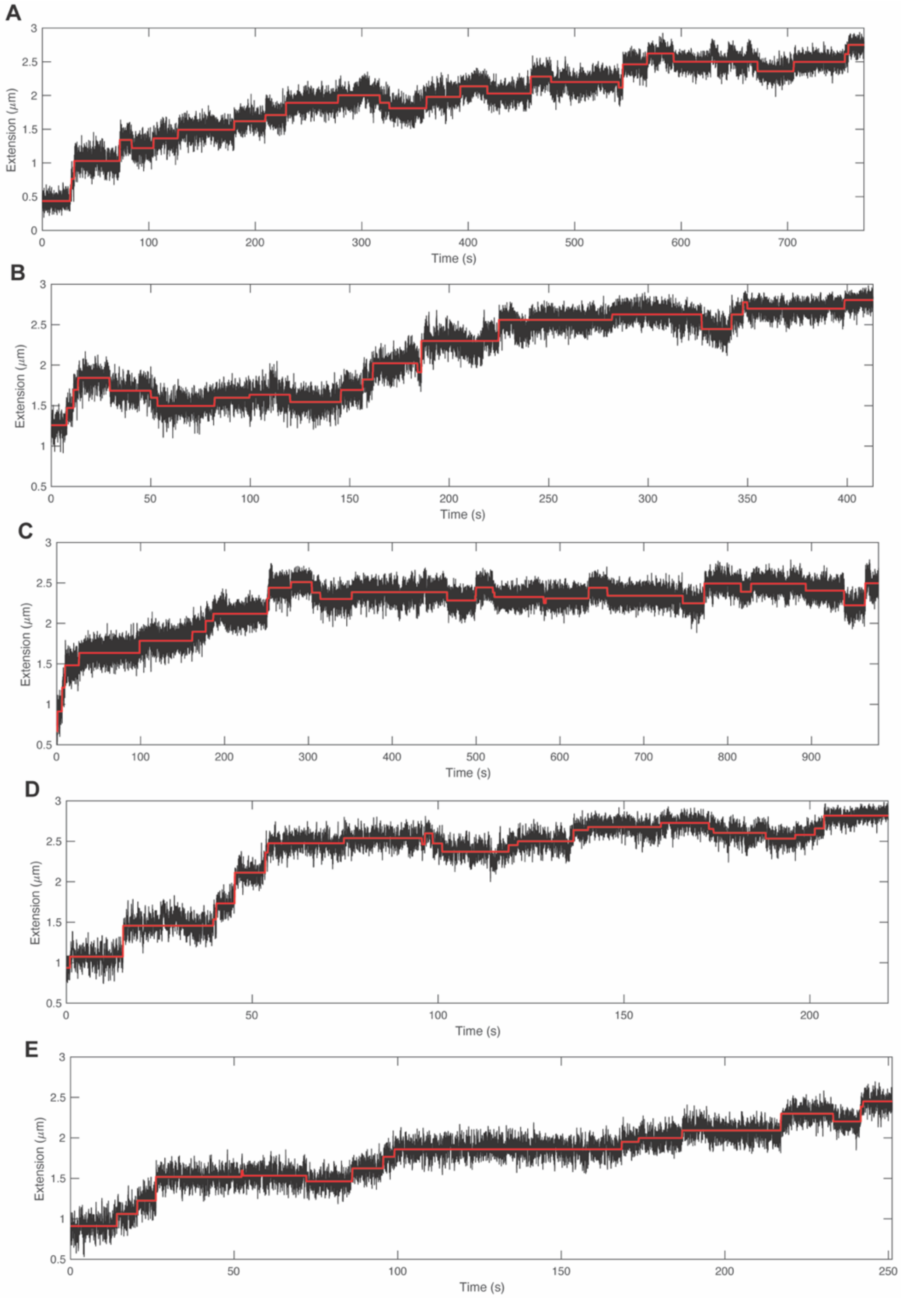


Figure S6. Representative Tel7dG strand invasion traces. Invasion traces show an increase in complexity when using the G-rich strand for invasion compared to the C-rich strand.

### Figure S7

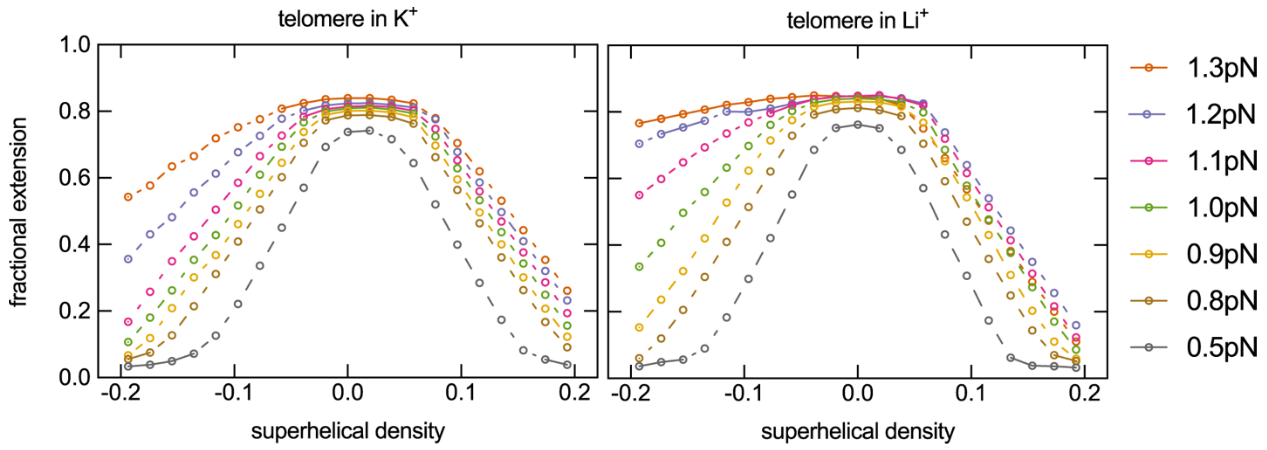


Figure S7. Telomere DNA torque response in $K^+$ (left) and $Li^+$ (right) at various forces.

**Figure S8**

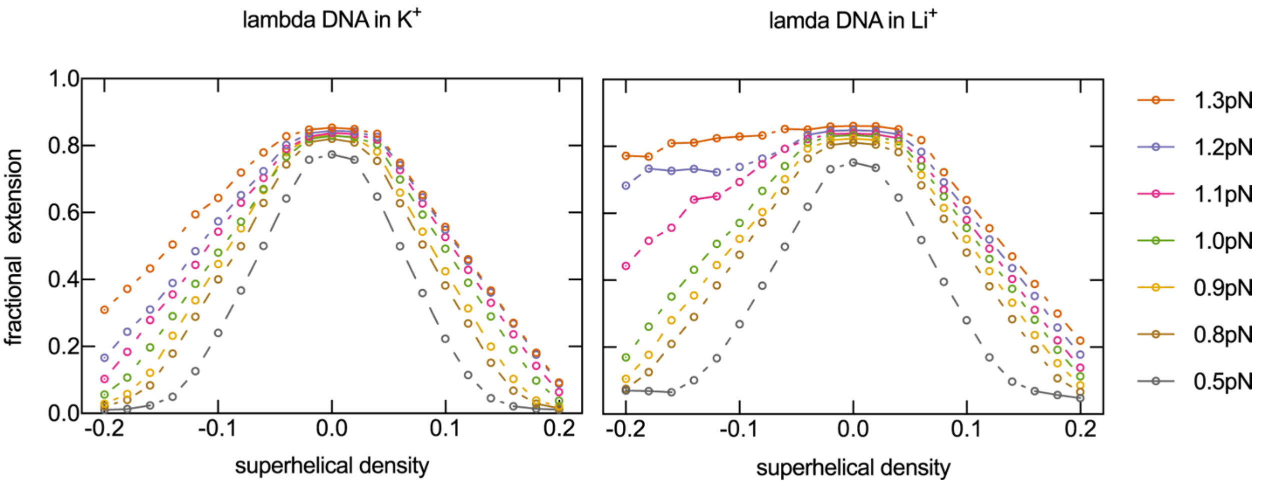


Figure S8. Lambda phage DNA torque response in $K^+$ (left) and $Li^+$ (right) at various forces.

**Figure S9**

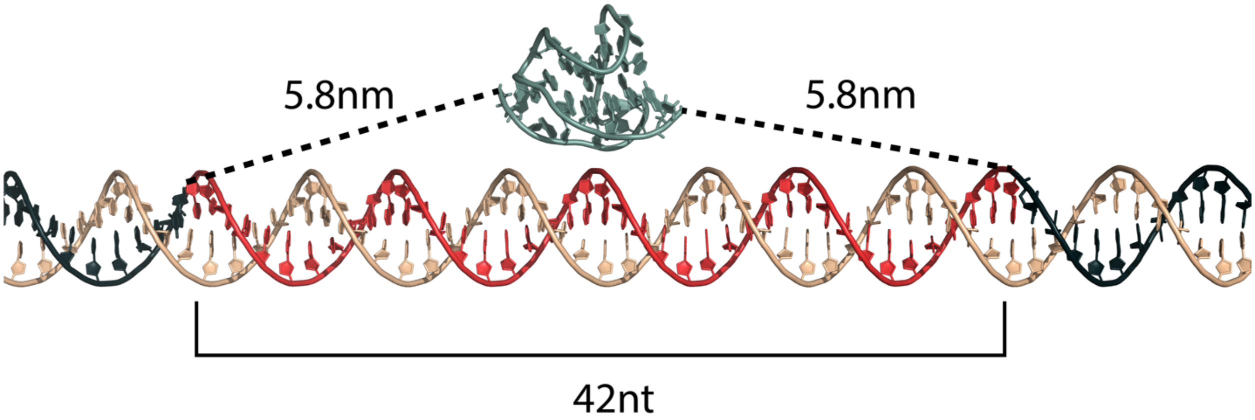


Figure S9**.** Model of a telomeric D-loop (TDL). The invaded 42 nt G-rich strand is modeled in red, a GQ modeled in green (PDB 2HY9). The measured distance between each end of the GQ and the sites of strand displacement in the TDL is 5.8 nm, consistent with the end extension length of ssDNA displaced but not folded into the GQ.

### REFERENCES


1. Stansel, R. M.; de Lange, T.; Griffith, J. D., T-loop assembly in vitro involves binding of TRF2 near the 3' telomeric overhang. *EMBO J* **2001,** *20* (19), 5532-40.
2. Long, X.; Parks, J. W.; Bagshaw, C. R.; Stone, M. D., Mechanical unfolding of human telomere G-quadruplex DNA probed by integrated fluorescence and magnetic tweezers spectroscopy. *Nucleic Acids Res* **2013,** *41* (4), 2746-55.
3. Lipfert, J.; Hao, X.; Dekker, N. H., Quantitative modeling and optimization of magnetic tweezers. *Biophys J* **2009,** *96* (12), 5040-9.
4. Yu, Z.; Dulin, D.; Cnossen, J.; Kober, M.; van Oene, M. M.; Ordu, O.; Berghuis, B. A.; Hensgens, T.; Lipfert, J.; Dekker, N. H., A force calibration standard for magnetic tweezers. *Rev Sci Instrum* **2014,** *85* (12), 123114.
5. Bustamante, C.; Marko, J. F.; Siggia, E. D.; Smith, S., Entropic elasticity of lambda-phage DNA. *Science* **1994,** *265* (5178), 1599-600.
6. Kerssemakers, J. W.; Munteanu, E. L.; Laan, L.; Noetzel, T. L.; Janson, M. E.; Dogterom, M., Assembly dynamics of microtubules at molecular resolution. *Nature* **2006,** *442* (7103), 709-12.
7. Woody, M. S.; Lewis, J. H.; Greenberg, M. J.; Goldman, Y. E.; Ostap, E. M., MEMLET: An Easy-to-Use Tool for Data Fitting and Model Comparison Using Maximum-Likelihood Estimation. *Biophys J* **2016,** *111* (2), 273-282.
